# Base Excision Repair Pathway Regulates Transcription-Replication Conflicts in Pancreatic Ductal Adenocarcinoma

**DOI:** 10.1101/2023.10.03.560510

**Authors:** Fan Meng, Anup K. Singh, Tiane Li, Marc Attiyeh, Fatemeh Kohram, Terence Williams, Yilun Liu, Mustafa Raoof

## Abstract

Oncogenic mutations (such as in KRAS) can dysregulate transcription and replication leading to transcription conflicts (TRCs). Unresolved TRCs can cause lethal DNA damage. Here, we sought to investigate the oncogene dependency of TRCs and TRC regulatory pathways in pancreatic ductal adenocarcinoma (PDAC). Human PDAC demonstrated 30-120-fold higher levels of TRC genomic signatures compared to breast, colon and lung cancer (p<0.001). TRCs were significantly enriched in human PDAC cells (Panc-1, BxPC3, MiaPaca2) compared to immortalized human pancreatic ductal epithelial (HPNE). Ectopic oncogenic KRAS(G12D) in HPNE cells enhanced TRCs, and TRC-related DNA:RNA hybrids (R-loops). Inhibition of KRAS or downstream effectors abrogated TRCs in Panc1 and MiaPaca2 cells. An siRNA screen identified several factors in the base-excision repair pathway as regulators of TRCs. In pharmacologic validation, inhibitors of APE1 endonuclease in BER pathway (Methoxyamine and CRT) enhanced TRCs. Mechanistic studies revealed that BER pathway inhibition severely altered RNA polymerase II dynamics at nascent DNA; causing RNAPII trapping and contributing to enhanced TRCs. The ensuing DNA damage activated Chk2-ATR pathway but not Chk1-ATM pathway. Co-treatment with ATR inhibitor (VX970) and BER inhibitor (methoxyamine) at clinically relevant doses, synergistically enhanced DNA damage and reduced cell proliferation in PDAC cells. The study uncovers a novel role of BER pathway defects and oxidative DNA damage in promoting TRCs. Our studies provide mechanistic insights into the regulation of TRCs in PDAC which has implications for genome instability and therapy in PDAC.

## INTRODUCTION

With a 5-year mortality rate of 90%, pancreatic ductal adenocarcinoma (PDAC) is among the most lethal human cancers (1). Not only is it highly fatal, PDAC is a high priority health problem; projected to be the second leading cause of cancer death by 2030. Therapeutic paradigms that have been successful in other solid tumors, have failed in PDAC. Therefore, innovative conceptual and therapeutic advances are urgently needed.

Oncogenic KRAS occurs in 95% of patients with PDAC and is a hallmark of cancer initiation and progression (2,3). Genetic ablation of mutant KRAS in animal PDAC models causes complete or near-complete tumor regression (4). This suggests that KRAS is a key therapeutic target in PDAC. However, targeting KRAS in human PDAC has been extremely challenging (5,6).

Protooncogene KRAS is a molecular switch that modulates signal transduction between membrane bound receptors and intra-cellular effector molecules to allow the cells to respond to environmental cues. Under quiescent conditions, KRAS is in a guanosine diphosphate (GDP)- bound inactive state which switches to a guanosine triphosphate (GTP)-bound activate state upon upstream signal transduction. Oncogenic mutations maintain KRAS in an activated state circumventing the need for upstream activation. In KRAS-mutant cancer cells, constitutive KRAS activity triggers RAF-MEK-ERK pathway activation (6,7). Downstream, ERK phosphorylation triggers further phosphorylation of ribosomal S6 kinase (RSK), serum response factor (SRF), E26 transformation-specific transcription factors (ETS) and ETS like-1 protein which promote enhanced transcription and protein synthesis of KRAS-activated genes. In parallel, GTP-KRAS also activates the PI3K-Akt-mTOR pathway causing profound effects on cell growth and survival through protein scavenging, metabolic rewiring, hyper-transcription, and enhanced protein synthesis (6,8,9). In addition to the increase in the level of transcription, oncogenic KRAS triggers aberrant enhancer activation which leads to dysregulated transcription (10).

Oncogenic mutations (such as in KRAS) can place significant demands on transcription and replication (2,11). In normal cells, transcription and replication are both spatially and temporally separated; whereas in oncogene-driven cancer cells there is loss of spatiotemporal separation (12). As a result of this dysregulation, the replication and transcription machinery frequently collide; causing transcription-replication conflicts (TRCs) at much higher rates in transformed cancer cells than in normal cells (12). Co-transcriptional processes that prevent stalling and backtracking of transcription machinery component such as RNA Polymerase II (RNAPII), can mitigate TRCs. For instance, RNAPII bound helicase RECQL5 can modulate transcription speed to prevent RNAPII stalling (13,14). However, persistent stalling of RNAPII can result in TRCs which are resolved by activation of transcription-coupled nucleotide excision repair pathway to remove the offending moiety on the DNA template followed by proteasome mediated degradation of RNAPII (15). Alternatively, RECQL5 can promote sumoylation of replication scaffold protein – proliferating cell nuclear antigen (PCNA) – to alter the histone composition at TRCs resulting dislodgement of RNAPII (16). In addition to stalled RNAPII, co-transcriptional DNA-RNA hybrids (R-loops) generated during transcription can also cause impediment to replication fork progression. Expectedly, RNA helicases that resolve R-loop such as Senataxin1 (SETX1), RNase H1 (RNH1), and DHX9 play a key role in TRC resolution pathways (17–19). Furthermore, certain DNA repair proteins serve non-canonical functions in TRC regulation. For instance, BRCA1 a homologous recombination pathway protein has been shown to interact with SETX1 to suppress R-loops and thereby mitigating TRCs (20).

Persistent stalling of replication machinery at TRCs can lead to DNA breaks which are resolved through downstream DNA repair pathways (21). TRCs occur more frequently in genomic areas that have long and difficult to transcribe genes – common fragile sites (CFS) – or areas of the genome that are both hyper transcribed and near replication origins – early replication fragile sites (ERFS) (22,23). These areas are particularly prone to copy number alterations and chromosomal rearrangements which are a characteristic feature of human PDAC. Unresolved TRCs cause lethal DNA damage (12,21,24). Thus, inhibiting TRC resolution and promoting TRC-induced DNA damage could be a promising therapeutic approach for KRAS-driven pancreatic cancer.

Despite the potential therapeutic significance of TRCs, the pathways that regulate TRCs in PDAC remain undefined. Here, we sought to characterize TRCs in PDAC and evaluate the causal role of oncogenic KRAS in promoting TRCs and DNA damage in PDAC. Further, we investigate DNA repair factors that mitigate TRCs. The study uncovers a novel role of BER pathway defects and oxidative DNA damage in promoting TRCs. We provide mechanistic insights into the regulation of TRCs in PDAC which has implications for genome instability and therapy in PDAC.

## METHODS

### Reagents and Antibodies

See reagents and antibody list (Supplemental Table)

### Cell Culture

HPNE E6/E7/st cells were obtained from a commercial source (ATCC, CRL-4036), and cultured with filtered medium (75% DMEM low glucose, 25% M3 base medium, 5% fetal bovine serum (FBS), 10ng/mL human recombinant EGF, 2mM L-glutamine, 750ng/mL puromycin). Constructed Doxy-inducible KRAS-G12D HPNE cells were cultured similarly except for replacing FBS to tetracycline-free FBS and supplemented with 10μg/mL Blasticidin (Fisher A11139-03). To express KRAS-G12D, 2 ng/mL Doxycycline was added to the medium at the indicated time in results. Panc-1, BxPC-3, Miapaca2, and HEK293 cells (CLS Cat# 300192/ p777_HEK293, RRID:CVCL_0045) were also commercially sourced (ATCC). Panc-1, Miapaca2, Hek293 were cultured with high-Glucose DMEM medium (Fisher, SH30423FS) supplemented with 10% FBS and 1% Pen-Strep (Sigma, 11074440001). The BxPC3 cell were cultured with RPMI1640 (Fisher, 11875119) supplemented with 10% FBS and 1% pen-strep.

### Generation of HPNE KRAS-G12D cell line

Tet-on inducible KRAS-G12D vector was generated using a standard cloning protocol. The KRAS-G12D cDNA were retrieved from pHAGE-KRAS-G12D vector by double digesting with Nhe-I and Bam-H1 followed by gel purification. The purified KRAS-G12D fragment was then ligated with mainframe of pCW-Cas9-Blast vector (addgene 83481) with Cas9 genes being replaced with KrasG12D. The sequence of recombinant pcw-KRAS-G12D-blast vector was verified with Sanger DNA Sequencing. To pcw-KRAS-G12D-blast vector was packaged into a viral assembly by co-transfecting with PMD2.G (addgene, 12259) and PSPAX2 (addgene,12260) into HEK293 T cells using polyethyleneimine (PEI) transfection reagents (Sigma 408727). The virus-containing medium was used to infect HPNE E6/E7/st40 cells with 8ug/ml polybrene. The infected cells were seeded at single cell density and subjected to antibiotic selection with full medium containing 10µg/ml blasticidin. Positively selected cells were expanded and KRAS-G12D expression was verified upon doxycycline induction by using western blot.

### Dot Blot Assay

To detect the levels of R-loops in parental and KRAS-G12D expressing clones, total genomic DNA was isolated from respective cells. Multiple serial dilutions of gDNA from each group starting from 500 ng were spotted on Hybond N+ positively charged nitrocellulose membrane designed to bind nucleic acids. After spotting the gDNA membrane was baked at 2 hours at 60 degree C crosslink the DNA on the membrane. Crosslinked membrane was blocked with 5% skimmed milk solution prepared in 1X TBS to minimize nonspecific protein-antibody interactions. Blocked membrane was incubated overnight with antibody against R-loops (Clone S9.6, Kerafast) under rotation. Next day, the membrane was washed thrice with 1X TBST before incubating in HRP-conjugated secondary antibody for 1 hour at room temperature. Corresponding level of R-loops was analyzed by using the chemiluminescence signal generated on membrane by using X-ray film.

### Western blot

A standard western blot protocol was used. Briefly, treated cells were lysed in RIPA buffer (10mM Tris-HCl Ph 8.0, 1mM EDTA, 0.5mM EGTA, 1% Triton-X 100, 0.1% Sodium Dexoycholate, 140mM NaCl) supplemented with protease and phosphatase inhibitor cocktail. After BCA assay quantification, cell lysates were mixed with equal volume of 2X Laemmli sampling buffer and loaded on 4-12% Tris-glycine gel for electrophoresis. The separated proteins were transferred onto PVDF membrane The target bands were sequentially probed by primary antibodies and secondary antibody as indicated in the specific results with dilutions ranging from 1:1000 to 1:5000. (The antibody source is indicated in the reagent/antibody list). The probed bands were revealed with West Pico plus luminol buffer (Fisher, 1863096)

### Isolation of protein in nascent DNA (iPOND)

We performed iPOND according to the previously published protocol (25). Briefly, cells were cultured in 15 cm plates to 60%-70% confluence. To pulse label the cells, EdU was added to final concentration 10 mM in cell medium for 10 mins at 37°C, followed by 10 mL 1% PFA fixing solution (20 min, room temperature. The PFA crosslink reaction was quenched by directly adding 1 mL of 1.3 M glycine buffer for 5 mins. The fixed cells were washed with DPBS at least twice and collected with cell lifter. The Cell pellet was retrieved by 900 g centrifuge for 10 mins and resuspended with 0.25% TritontX100-DPBS for permeabilization at room temperature for 30 mins, then washed with 0.5% BSA DPBS and DPBS buffer. Cells were then centrifuged at 900 g to retrieve the cell pellet. Click reaction was performed to conjugate EdU and biotin by resuspending and mixing the cell pellet in click solution (1.76 mg/mL sodium L-ascorbate; 2 mg/mL CuSO_4_; and 10μM Biotin azide) for 1 hour at room temperature. For negative control “no pulse” sample, the click solution excluded Biotin azide. The clicked samples were then washed with BSA and DPBS at least one time each, and resuspended with 2% SDS lysis buffer (50mM Tris-HCl pH=8 with 1μg/mL Leu-peptin and aproptinin) on ice for 10 mins, followed by sonification (Bioruptor Pico, Diagenode). The sonicated samples were centrifuged 16,100 g for 10 mins at room temperature, and 10μL supernatant mixed with Laemmli buffer was used as input sample. Whole supernatant was 1:1 diluted with DPBS supplemented with Leueptin and aproptinin as previous indicated, then mixed with 50μL pre-washed Streptavidin-agarose resins under gentle rotational mixing at 4°C overnight to pull-down biotin conjugated nascent DNA protein complexes. After streptavidin-biotin conjugation, the beads were washed several times to wash away non-specific binding proteins. The streptavidin bound protein complexes were then retrieved and subjected to western blot as above to probe for target proteins.

### Cell fractionation and protein purification

Cell fractionation was performed as described previously with some modifications (16). Briefly, treated cells were trypsinized and washed one time with DPBS. The cell pellet weight was measured and lysed with 3 volumes (3μL/mg cell pellet) of cytoplasmic buffer for 30 mins (10⍰mM Tris–HCl pH 7.5, 0.34⍰M sucrose, 3⍰mM CaCl2, 2⍰mM MgCl_2_, 0.1⍰mM EDTA, 1⍰mM DTT, 0.5% NP40, 40⍰mM NEM) supplemented with protease and phosphatase inhibitors. The whole lysate was then centrifuged at 2400⍰×⍰g for 5 mins to collect the nuclear pellet, which was resuspended in nuclear buffer (20⍰mM HEPES pH 7.5, 1.5⍰mM MgCl2, 1⍰mM EDTA, 150⍰mM KCl, 0.1% NP40, 1⍰mM DTT, 10% Glycerol) at 1.5 times the volume of pellet, followed by gentle homogenization with a 21G needle. After homogenization, nuclear lysates were then centrifuged with 18,000⍰×⍰g for 30⍰min to obtain the intact chromatin pellet. For separating CB:RNA+ and CB:RNA− fraction, the chromatin pellet was first incubated with RNase A in RNase A buffer (50⍰mM Tris–HCl pH 8.0, 10⍰mM EDTA, 150⍰mM NaCl, RNase A 10⍰µg/mL) at 1.5 times the volume of pellet, overnight (under gentle rotation at 4°C). The solubilized protein in the supernatant was collected as the CB:RNA+ fraction. The remaining pellet was then digested with benzonase (overnight at 4°C) in nuclease buffer(,(20⍰mM HEPES pH 7.5, 1.5⍰mM MgCl2, 1⍰mM EDTA, 150⍰mM KCl, 10% Glycerol, 0.5⍰U/µL benzonase) at 1.5 times the volume of pellet. The whole lysates were then mixed (50:50) with 2 x Laemmli sample buffer and were boiled for 20 mins. Resulting sample was then collected as the CB:RNA-fraction. The CB:RNA+ and CB:RNA-were then subjected to western blotting.

### PLA assays, flowcytometry and immunocytochemistry

For high-throughput experiment, cells were seeded on collagen-coated 96 well plate. Whereas for validation experiments, cells were cultured on collagen-coated glass on 6 well plate. After reaching ∼60% to 70% confluence, the cells were fixed with 4% Paraformaldehyde (PFA) for 15 mins at room temperature. Fixed cell was washed with DPBS twice, and then permeabilized by adding 0.5% NP40 on ice for 5 mins. Subsequently, the cells were then washed with DPBS 3 times and then blocked with 2% BSA-DPBS buffer at room temperature for 1 hour. The cells were then incubated with RNAPII CTDS2 (1:500) and PCNA (1:500) antibodies at 4°C overnight. Proximity ligation was then performed using manufacturer instructions supplied with the kit (Sigma Aldrich). The samples were stained with DAPI and images were acquired using either Olympus Ix71 inverted fluorescence tissue microscope (single experiment) or ZEISS LSM900 confocal light microscopy (high-throughput experiment). The images were then processed and analyzed with QuPath and Image J (RRID:SCR_003070) to calculate the PLA mean intensity per nucleus in each captured image.

For flowcytometry PLA assay, treated cells with 60-70% confluence were trypsinized, washed at least one time with DPBS, then fixed with 4% PFA for 15 mins followed by permeabilization with 5 minute incubation with 0.5% NP-40. The fixed cell pellets were then collected after centrifugation at 400xg for 5 mins and washed with DPBS twice, blocked with commercial block buffer (supplied in kit) at room temperature for 1 hour, and then incubated with RNAPII-CTD S2 (1:100) and PCNA (1:100) antibody overnight in 4°C. PLA reactions were performed using the manufacturer supplied instructions in the kit (Sigma Aldrich). PLA-fluorescent labelled cells were passed through a 100µm strainer, and the PLA signal was measured using Attune NXT cytometer. FlowJo (RRID:SCR_008520) for data processing, analysis and visualization.

### Cell proliferation with Real-time cell analysis (RTCA)

For measuring the inhibitory effect of methoxyamine (MX) and ATR inhibitor (VX970) in Panc1 and MIA Paca2 cells, the RTCA assay was used to measure real-time cell growth. Approximately 4000 cells were seeded on an E-plate (Agilent 05469830001). Cell growth curves were recorded on xCELLigence RTCA DP (Agilent, CA, USA) at 15-min intervals under standard cell culture conditions (37°C, 5% CO2). Indicated treatments were added 24 hours after plating.

### TRC mutational signature analysis

The International Cancer Genome Consortium (ICGC) was queried to retrieve whole genome sequencing and RNA expression data pertaining to breast (project BRCA-US), lung (project LUAD-US), colorectal (project COAD-US), and pancreatic (projects PACA-AU and PACA-CA) tumor types. The dataset exclusively included tissue samples from primary tumors, and patients had not undergone any prior treatment. Custom software, implemented in the Java programming language, was employed to quantify the frequency of mutations in the promoter or promoter-proximal (PPP) regions, PPP regions associated with highly transcribed genes (PPP-HTG), and common fragile sites (CFS). The definition of PPP regions covered a region spanning one kilobase upstream of transcription start sites (TSS), relying on a predefined set of regions acquired from the UCSC Genome Browser (26). PPP-HTG regions were identified as regions located one kilobase upstream of genes exhibiting expression levels at or above the 75th percentile for the respective sample. As an aside, each donor in the ICGC database can contain one or more specimens, and multiple samples from each specimen can be used for analysis. To ensure compatibility and consistency in the results, only mutation and expression data from the same specimen (either from one sample or multiple samples) were employed for analysis. CFS regions were defined based on pre-existing data from published sources (27), and subsequently cross-referenced with cytoband start and end positions acquired from the UCSC Genome Browser (28).For each tumor type, the median number of mutations and the corresponding interquartile ranges were calculated.

### Statistical Analysis

Each experiment was independently repeated at least twice, and representative results are shown. For individual experiments at least 3 technical replicates were analyzed unless stated otherwise in the results section. For all in vitro experiments, treatment groups were randomly assigned. Image acquisition for in vitro experiment was automated to reduce bias and ensure blinding when possible. Statistical analyses were performed using Prism 10.0 (GraphPad Prism, RRID:SCR_002798). Data are presented as the mean ± SD, unless otherwise indicated. Differences between the two different groups were evaluated using a paired or un-paired student t-test for parametric data. For non-parametric data, indicated non-parametric tests were used. P < 0.05 was defined as statistical significance. Tukey test was used for multiple comparisons after one-way ANOVA. Formal power analysis was not performed.

## RESULTS

### Human PDAC demonstrates high levels of TRCs

To determine the extent to which TRCs play a role in human PDAC, we analyzed gene expression profiles of a panel of known TRC proteins (13,17–20) in tumor and matched normal tissue using TCGA and GETX datasets. The results in Figure 1A demonstrate a statistically significant increase in gene expression of proteins that are known to resolve TRCs (p<0.001). Similarly, expression of these proteins was uniformly elevated in PDAC compared to other tumor types (Figure S1). We then evaluated DNA mutational signatures associated with TRC across common human solid tumors (breast, non-small cell lung, colorectal and pancreas) using data from the International Cancer Genome Consortium. Promoter and promoter-proximal mutations in highly transcribed genes[10] as well as common fragile site (CFS) mutations are associated with TRCs (21). As shown in Figure 1B these mutations were much more prevalent (30-120-fold) in PDAC in comparison to other common solid tumors. We then evaluated TRCs in human pancreatic cancer cells (Panc1) vs. human pancreatic normal epithelial cells (HPNE). We isolated proteins on nascent DNA (iPOND) using a well-established technique (Figure 1C) (25). HPNE or Panc1 cells were pulsed with 5-Ethynyl-21-deoxyuridine (EdU) to label nascent DNA and compared to non-pulsed cells. The EdU was clicked to biotin-azide and protein complex in association with biotin-EdU-labelled DNA were isolated using streptavidin pull-down. The isolated protein complexes were examined for transcription protein, RNAPII and replication factor, PCNA. We noted that RNAPII is significantly enriched at the replication fork in Panc1 cells compared to HPNE cells, whereas the levels of PCNA were similar. This finding demonstrates high levels of TRCs in Panc1 cells compared to HPNE cells.

**Figure 1.**
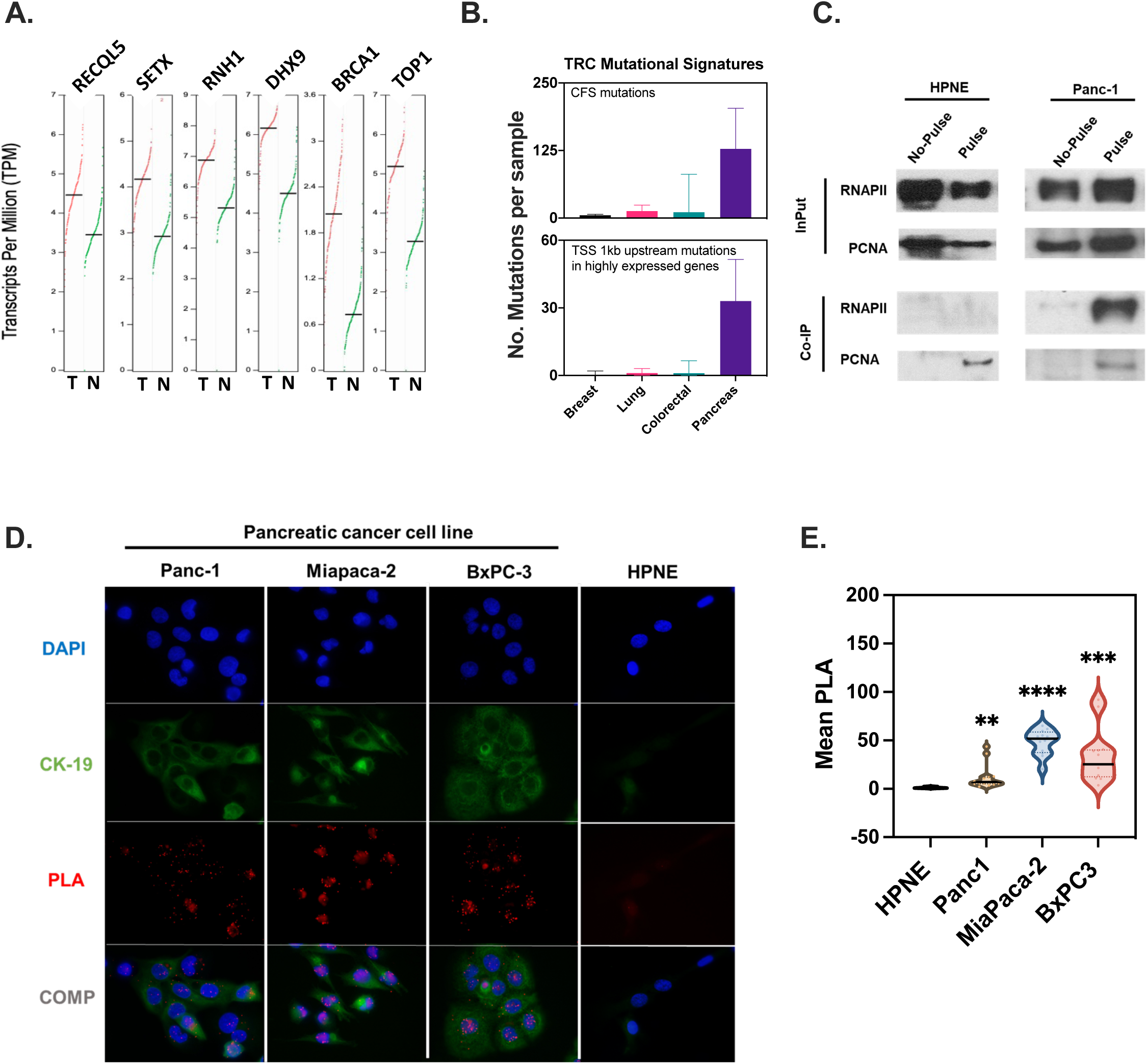
Human PDAC demonstrates high levels of TRCs. **A Gene-expression analysis** of known proteins involved in TRC resolution pathways are upregulated in PDAC tumors (**T**) vs normal tissue (**N**) (All T vs. N comparisons – Student t-test, p-value <0.001). Tumor data source TCGA project, Normal data source is GETx project. **B. TRC mutational signatures** significantly enriched in human PDAC vs. other solid tumors. Median mutations per tumor sample are shown. Error bars indicate interquartile range. TSS – transcription start site; highly transcribed (>75^th^ percentile for gene expression). (P<0.001, PDAC vs. other tumor types, Mann-Whitney U test) **C. Isolation of Proteins on Nascent DNA.** (iPOND). Newly synthesized DNA was labelled with EdU in pulsed HPNE and Panc1 cells for 2 minutes. EdU was conjugated to biotin and the proteins on EdU-biotin labeled DNA were pulled down using streptavidin beads (Co-IP). Representative western blot demonstrates presence of RNAPII at replication forks in Panc1 pulsed cells but not in non-pulsed Panc1 cells, or pulsed/ non-pulsed HPNE cells; consistent with the high burden of TRCs in Panc1 cells but not in HPNE cells. **D & E. Immunocytochemistry PLA assay (PCNA-RNAPII)** demonstrates high TRCs in all human PDAC cell lines compared to HPNE cells. (Representative experiment from a duplicate is shown. Welch’s unpaired t-test, each cell line was compared to HPNE cell line, *P=0.0062, **P=0.0009, ***P<0.0001)

To confirm this finding, we measured TRC levels in a panel of human pancreatic cancer cell lines using PLA assay. PLA is an established, highly sensitive and specific immunohistochemical tool for detecting two proteins within 40 nm of each other (29). We adapted and optimized this method to detect the RNAPII-PCNA proximity as a marker of TRCs. We chose these two proteins as RNAPII is a central component of transcription machinery and PCNA is a hub within the replication machinery. We found elevated levels of TRCs in these cell lines compared to the HPNE control cell line which did not show any detectable TRCs (Figures 1D, E). Interestingly, the elevated levels of TRCs were not unique to KRAS mutated cell lines as KRAS-wildtype BxPC3 cells also demonstrated high levels of TRCs. BxPC3 cells have a downstream BRAF activating mutation that may phenocopy KRAS mutations with regards to TRCs. Collectively, these findings demonstrate that human PDAC demonstrates high levels of TRCs.

### Oncogenic KRAS causes TRCs and DNA damage

To determine the impact of oncogenic KRAS(G12D) on TRCs we generated a doxycycline inducible system for KRAS(G12D) expression. We used commercially sourced telomerase-immortalized human pancreatic ductal-derived (HPNE) cell line (as previously published (30)). This cell line expresses human papillomavirus E6 and E7 oncogenes, which block the function of the p53 and Rb tumor suppressors, respectively. In addition, SV40 small t antigen is expressed as it is required to allow oncogenic transformation upon KRAS (G12D) expression. A doxycycline inducible KRAS(G12D) vector was stably transfected in the HPNE cells, and its expression was verified by western blot. As shown in Figure 2A-B, upon doxycycline induction, KRAS(G12D) expression increased over time. Further, KRAS(G12D) induction caused downstream oncogenic signaling as manifested by increased Akt phosphorylation. This resulted in replication stress and DNA damage as evidenced by increase γH2AX phosphorylation over time.

**Figure 2.**
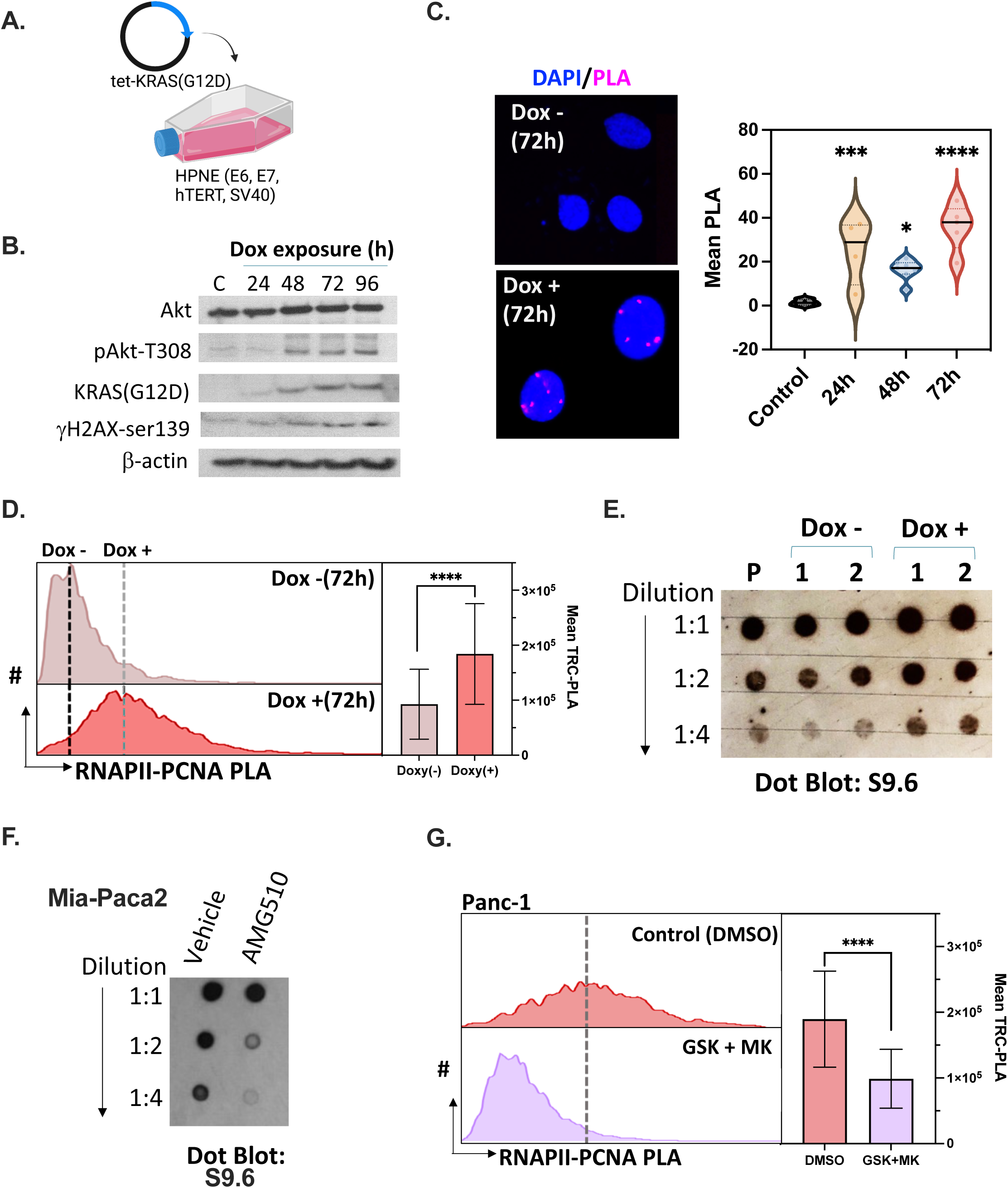
Oncogenic KRAS causes TRCs and DNA damage. **A. Experimental set-up.** HPNE cells were stably transfected to express KRAS(G12D) oncogene upon doxycycline induction. **B.** Representative **western blot** of the indicated proteins is shown from whole cell lystes of HPNE cells indicating downstream activation of Akt pathway and DNA damage upon KRAS induction. **C. Immunocytochemistry** and **D. Flowcytometry** demonstrates accumulation of TRCs upon KRAS induction in HPNE cells over time as assessed by PCNA-RNAPII proximity ligation assay. (Immunocytochemistry, Violin plots are shown. Dunnett’s multiple comparison test: control vs. 24h **p=0.0043; control vs. 48h *p=0.0397, control vs. 72h ****p<0.0001). (Flowcytometry, Mean and Standard deviation is shown, unpaired t-test, ****p<0.0001) **E. Dot blot assay.** HPNE parental cells (P), and two clones (1 & 2) stably transfected with doxycycline-inducible KRAS(G12D) oncogene were assessed for DNA-RNA hybrids (R-loops) with or without KRAS(G12D) induction for 3 weeks. Equal amount of Genomic DNA was blotted and probed with R-loop specific antibody (S9.6). **F. Dot blot assay.** Mia-Paca2 cells were treated with KRAS(G12C) inhibitor - Sotorasib (AMG510, at 6.5nM) or vehicle for 24 hours. Genomic DNA was extracted, and equal amount was blotted and probed with R-loop specific antibody (S9.6). Representative dot blot is shown. **G. Flowcytometry assay.** Panc-1 cells were treated with MEK inhibitor (GSK1120212, GSK) and AKT inhibitor (MK2206, MK) or vehicle control at sub-lethal dosage for 24 hours and TRCs were measured using flowcytometry bases RNAPII-PCNA PLA assay. (Mean and Standard deviation is shown from a representative experiment is shown, unpaired t-test, ****p<0.0001)

We then examined the extent to which oncogenic induction increased TRCs. This was evaluated using a proximity ligation assay (PLA). As shown in Figure 2C-D, KRAS(G12D) induction in HPNE cells by administration of doxycycline caused increased TRCs compared to control conditions. TRCs characteristically generate unusual non-B DNA structures such as DNA-RNA hybrids (R-loops) (31). To test if KRAS(G12D) induction in HPNE cells increases R-loops, parental clone and two different KRAS(G12D) expressing clones were assessed for R-loops using dot-blot with or without 3 weeks of doxycycline exposure. We used an R-loop specific antibody (S9.6) to detect R-loops in genomic DNA isolated from HPNE cells. The results shown in Figure 2E and S2A demonstrate increase in R-loops upon KRAS(G12D) induction by doxycycline in both clones; whereas, cells without doxycycline exposure had R-loop levels comparable to parental HPNE cell line. To determine if the observed R-loops in PDAC are generalizable to other KRAS mutations, we exposed MiaPaca2 cells harboring KRAS(G12C) mutation to a specific KRAS(G12C) inhibitor – AMG510 (Figure 2F and S2B). The findings demonstrated that inhibition of KRAS(G12C) mitigated R-loops at clinically relevant concentrations. Since KRAS(G12D) inhibitors were not available to us at the time of experiments, we treated Panc1 cells with MEK inhibitor (GSK1120212, GSK) and/ or AKT inhibitor (MK2206, MK) or vehicle control at sub-lethal dosage for 24 hours and measured TRCs using flowcytometry based RNAPII-PCNA PLA assay. We found that complete abrogation of KRAS(G12D) downstream signaling also completely abrogated TRCs in Panc-1 cells (Figure 2G and S2C). Collectively, these data demonstrate the causal role of oncogenic KRAS in propagating TRCs in PDAC.

### Base excision repair (BER) pathway regulates TRCs in PDAC cells

Our data suggests that a high level of TRCs cause sub-lethal genome instability upon oncogenic KRAS induction (as evidenced by enhanced γH2AX, Figure 2B). We hypothesized that established PDAC cells must require dependence on TRC resolution factors to maintain fitness as an adaptive response. To identify proteins functionally involved in the regulation of TRCs in human PDAC Panc1 cells, we conducted an siRNA knock down screen using a library targeting 200 DNA damage response factors (Dharmacon, Figure 3A). After 72 hours of siRNA exposure, TRCs were measured using a high throughput RNAPII-PCNA PLA assay. As shown (Figure 3) the screen identified several known TRC-resolution factors including: RECQL5 (13,14,16), TRIM28 (32), SETX (17), TOP2A (21), and transcription-coupled nucleotide excision repair proteins (ERCC6/8)(33). Importantly, this screen identified several proteins with no prior known involvement in TRC prevention or resolution. Interestingly, many of the top hits are proteins with established roles in BER pathway including MPG, POLB, APTX, NEIL2, SOD1, FEN1, PARP1, and APEX1 (Figure 3B)(34). Multiple hits in the same pathway strongly implicate BER in mitigating TRCs (Figure 3C).

**Figure 3.**
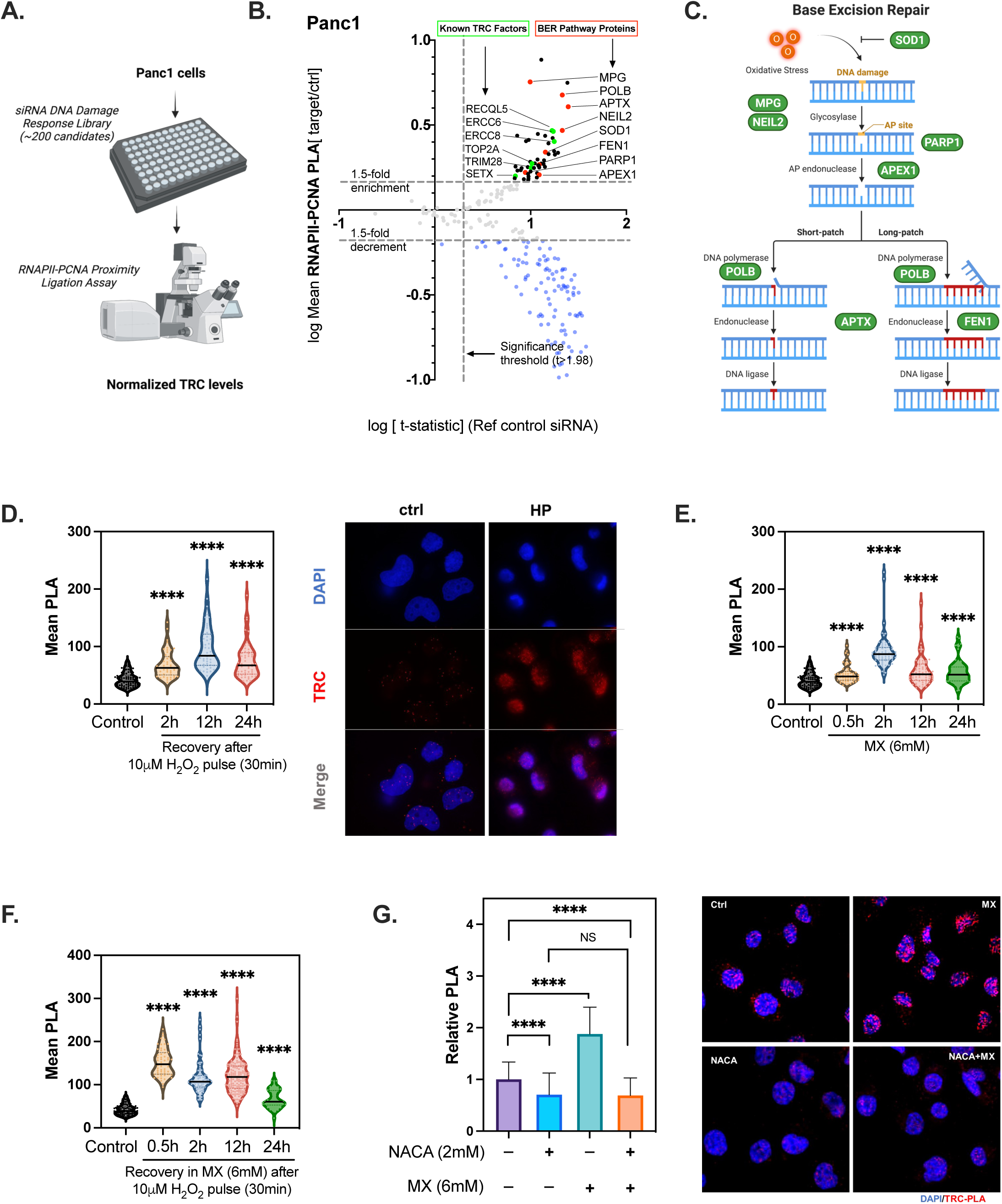
BER pathway regulates TRCs in PDAC cells. **A.** Experimental set-up of siRNA screen. **B.** Unbiased siRNA screen to identify TRC regulators in Panc1 cells. Sub-confluent cells were cultured in 96-well plates and exposed to siRNA for 72 hours. TRCs were quantified by measuring mean RNAPII-PCNA PLA signal intensity per well. The data were normalized to number of cells and RNAPII signal intensity per well. At least 25 cells were quantified per well. X-axis represents log transformation of fold enrichment/ decrement over control condition. Y-axis represents log of t-statistic. t>1.98 signifies TRC levels significantly different than control condition. **C.** Schematic of base excision repair pathway demonstrating the location of “hits” from the siRNA screen within the pathway. **D-G. Immunocytochemistry.** RNAPII-PCNA PLA was performed to quantify TRCs after indicated treatment conditions. Representative experiments are shown. (****p<0.0001, Welch’s unpaired t-test vs. control or otherwise indicated comparisons). MX=methoxyamine. NACA=N-Acetyl Cysteine Amide.

The BER pathway is very important for repairing oxidative DNA damage (34). Thus, we hypothesized that oxidative damage and the BER pathway might be contributing to and helping to resolve TRCs respectively. TRC associated BER protein expression was significantly higher in tumors compared to normal (Figure S3A). In pan-cancer analysis most BER proteins (except for FEN1 and PARP1) were not associated with poor survival (Figure S3B). However, in PDAC gene expression of several TRC-associated BER proteins was associated with poor survival including FEN1, MPG, and APEX1 (Figure S3B and S3C).

To validate the role of oxidative DNA damage in regulating TRCs, we manipulated oxidative DNA damage pharmacologically. Panc1 cells were exposed to a brief 30 min sub-lethal pulse of H_2_O_2_ (10μM) and TRCs were analyzed using RNAPII-PCNA PLA assay over 24 hours (Figure 3D). This experiment demonstrated a brisk increase in TRCs at 2 hours, with peak at 12 hours followed by a slight decline at 24 hours. Since H_2_O_2_ can cause widespread non-specific oxidative damage, we asked if upregulation of TRCs can be attributed to oxidative DNA damage. To test this hypothesis, Panc1 cells were exposed to sub-lethal concentrations of methoxyamine (MX - CH_3_ONH_2_). MX is a specific covalent inhibitor that binds to apurinic/ apyrimidinic (AP) sites that result from processing of oxidative DNA damage by BER pathway glycosylases (35). MX treatment significantly enhanced TRCs in a manner similar to H_2_O_2_ exposure (Figure 3E), with a combination of MX and H_2_O_2_ causing additional increase in TRCs (Figure 3F). Similar observations were noted with APE1 inhibitor - CRT0044876 (Supplemental Figure S3D and E)(36). The increase in TRCs noted in Panc 1 cells with MX was completely abrogated below baseline levels by exposure to anti-oxidant – N-Acetylcysteine Amide (NACA) (Figure 3G). Interestingly, NACA also mitigated TRCs in Panc1 cells at baseline suggesting oxidative DNA damage may contribute to basal level of TRCs in PDAC cells (Figure 3G).

Collectively, these findings highlight a novel role of BER pathway in regulating transcription-dependent replication stress in PDAC cells.

### BER inhibition impacts RNAPII dynamics at TRCs

To characterize the mechanism by which BER inhibition enhances TRCs, we evaluated RNAPII dynamics at nascent DNA after BER inhibition. We hypothesized that AP sites created by oxidative DNA damage would stall RNAPII on template strand and would enhance the probability of TRCs. To test this hypothesis, we isolated protein complexes on nascent DNA via iPOND from Panc1 cells, with and without exposure to MX for a varying duration. The complexes were then subjected to western blotting to identify the presence of transcription, replication, and DNA damage marker proteins. As shown in Figure 4A and S4A, there was enrichment of transcription and replication complexes in pulsed compared to non-pulse control. The MX treatment significantly enhanced the association of PCNA, RNAPII catalytic sub-unit (RPB1). Over time RPB1 and PCNA stayed associated with the nascent DNA, however, the RNAPII became dephosphorylated as evidenced by declining levels of Ser2 phosphorylation of RNAPII C-terminal domain (RNAPII-CTDS2). Furthermore, the decline in RNAPII-CTDS2 was anti-correlated with the rising levels of γH2AX indicating DNA damage at nascent DNA.

**Figure 4.**
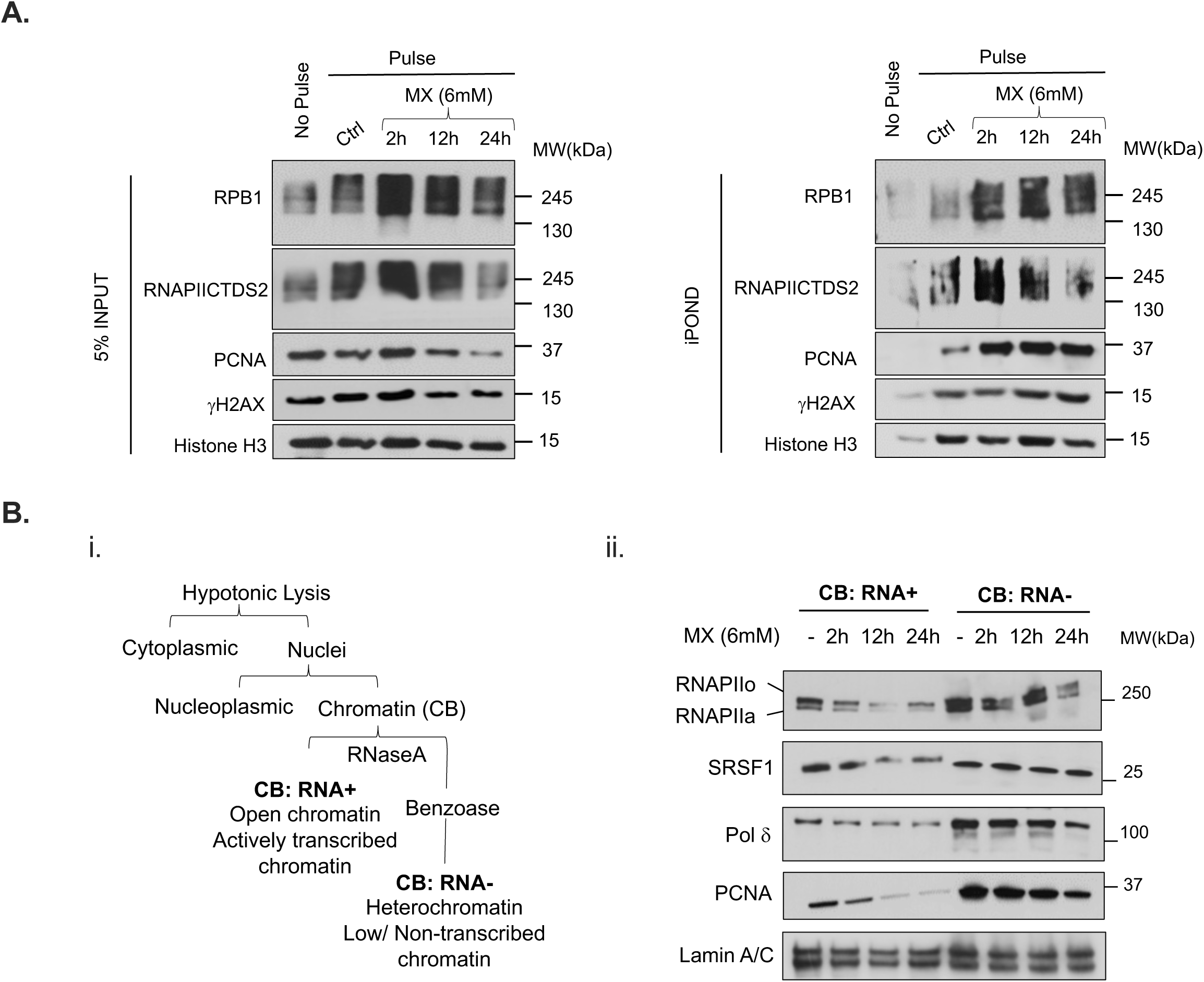
BER inhibition impacts RNAPII dynamics at TRCs. **A. Isolation of Proteins on Nascent DNA.** (iPOND). Newly synthesized DNA was labelled with EdU in pulsed Panc1 cells for 2 minutes after indicated treatment conditions. EdU was conjugated to biotin and the proteins on EdU-biotin labeled DNA were pulled down using streptavidin beads (Co-IP). Non-pulsed cells were used as an additional control. Representative western blot demonstrates presence of catalytically active sub-unit of RNAPII (RPB1), elongating RNAPII (RNAPIICTDS2), replication machinery component (PCNA) and DNA damage marker (γH2AX). Histone H3 is used as a loading control. Methoxyamine (MX) treatment significantly enhanced both replication and transcription machinery components at replication forks as well as DNA damage. The elongating RNAPII (CTDS2) decreased overtime suggesting stalling of RNAPII at TRCs. **B. i.** Schematic diagram of the cell fractionation to enrich proteins associated with transcriptionally active open chromatin structure (CB:RNA+) from proteins that are bound to heterochromatin, or from regions not highly transcribed (CB:RNA−). ii. Representative western blot analysis of total RNAPII (o=active or, a=inactive fraction), splicing factor SRSF1, DNA polymerase (Pol δ) and replication scaffold (PCNA) in CB:RNA+ and CB:RNA− fractions prepared from Panc 1 cells after indicated treatments.

To determine if stalling and inactivation of RNAPII in response to BER inhibition is preferentially at sites of actively transcribed chromatin, we isolated protein complexes at these sites from Panc1 cells. We used a previously established chromatin fractionation protocol that enriches for protein complexes at transcriptionally active chromatin (CB:RNA+) and compared these to protein complexes at non-enriched chromatin (CB: RNA-) (Figure 4Bi). This experiment demonstrated that MX treatment caused a rapid and robust decrease in levels of RNAPII (both, active – RNAPIIo and inactive – RNAPIIa) at CB:RNA+ fraction but not to the same extent in CB:RNA-fraction. Similarly, we noted decrease in levels of splicing factor - SRSF1 and replication machinery components (PCNA and DNA polymerase δ) at actively transcribed chromatin but Lamin A/C (loading control (37)) which associates with both transcribed and non-transcribed chromatin remained unchanged (Figure 4Bii and S4C).

Collectively, these findings suggest that BER inhibition causes replication and transcription stress predominantly at transcriptionally active chromatin.

### BER inhibition causes transcription-dependent DNA damage and activates ATR-Chk1 pathway

To determine the consequence of replication and transcription stress by MX at transcriptionally active chromatin, we evaluated DNA damage marker (γH2AX) with or without co-treatment with transcription inhibitor (DRB 6-dichloro-1-beta-D-ribofuranosylbenzimidazole). As demonstrated in Fig 5A, consistent with prior results, MX treatment resulted in decreased levels of both non-phosphorylated (RNAPIIa) and phosphorylated forms of RNAPII (RNAPIIo and RNAPII CTDS5) in whole cell extracts of Panc1 cells. In contrast, DRB treatment for 2 hours caused swift inhibition of RNAPII phosphorylation. The DNA damage increased with MX exposure but was significantly abrogated by DRB co-treatment at the 24h timepoint. These results demonstrate that DNA damage resulting from BER inhibition in PDAC cells is at least partly transcription dependent.

**Figure 5.**
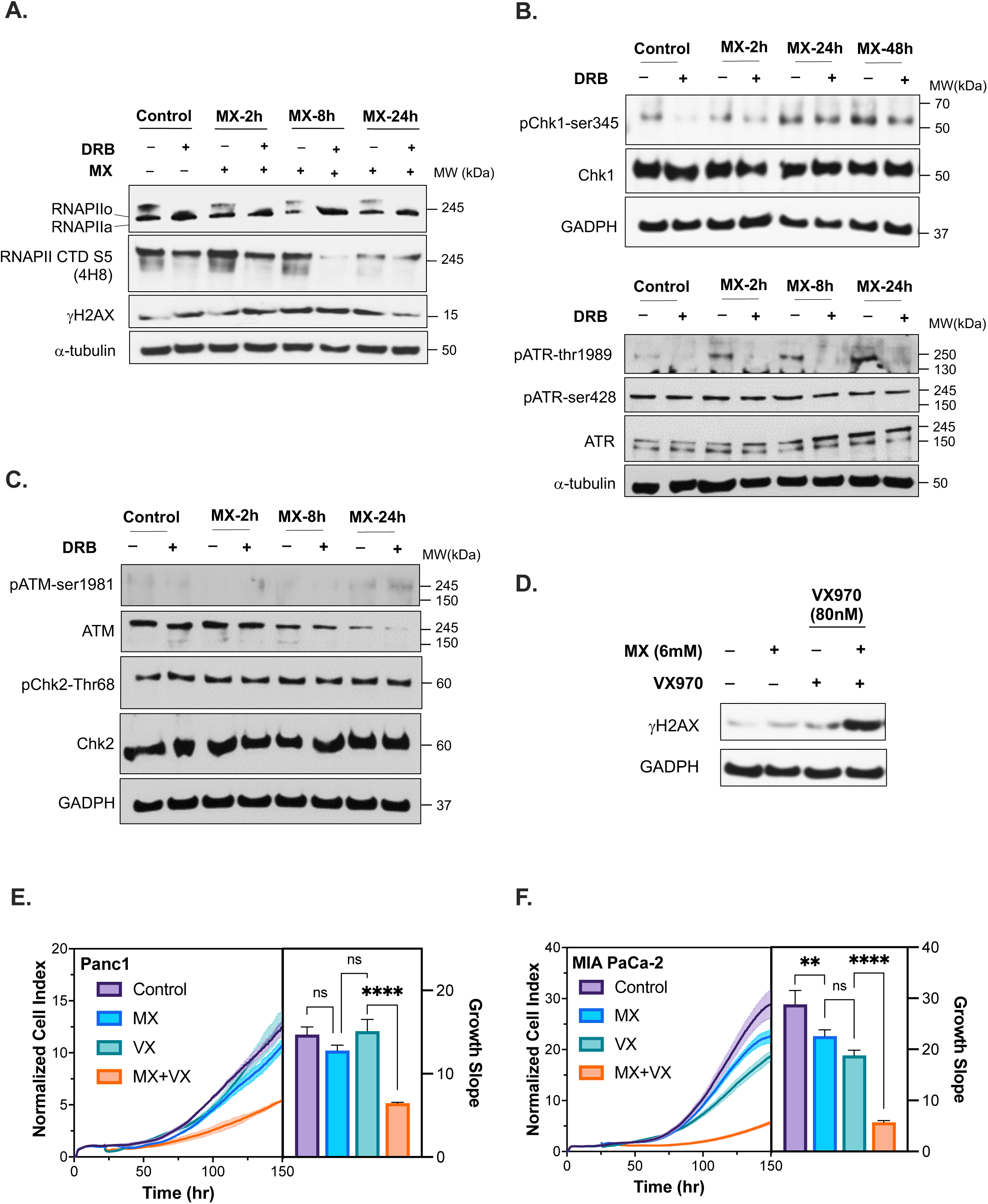
BER inhibition causes transcription-dependent DNA damage and activates ATR-Chk1 pathway. **A.** Representative Western Blot analysis of RNAPII (o=active or, a=inactive fraction), early elongating RNAPII (RPB1 CTDS5), and DNA damage marker (γH2AX) in Panc1 whole-cell extracts prepared from control, MX or DRB-treated cells as indicated. DRB is a transcription inhibitor and was administered for the last 2h of the experiment at 100μM. α-tubulin was used as a loading control **B.** Representative Western Blot analysis of ATR-Chk1 pathway in Panc1 whole-cell extracts prepared from control, MX or DRB-treated cells as indicated. α-tubulin and GADPH were used as loading controls. **C.** Representative Western Blot analysis of ATM-Chk2 pathway in Panc1 whole-cell extracts prepared from control, MX or DRB-treated cells as indicated. GADPH was used as a loading control. **D.** Representative Western Blot analysis of DNA damage (γH2AX) after ATR pathway inhibition using ATR inhibitor (VX970) in Panc1 whole cell extracts co-treated with or without MX as indicated. GADPH was used as a loading control. **E-F.** Real-time cell analysis of Panc1 or Mia-Paca2 cells treated with or without MX (12mM), VX970 (160nM) or combination as shown. Cell-index is a unitless measure of cellular impedance of electron flow caused by adherent cells depicting cell growth and confluence. Lines indicate average of three replicates and error bars indicate standard deviation. (Slope comparison: One-way ANOVA with Tukey multiple comparison test: *p<0.034, **p<0.0021, ***p<0.00021, ****<0.0001)

DNA damage from TRCs can activate downstream DNA damage checkpoint pathways. Hamperl et. al. have previously demonstrated that head-on TRCs, increase R-loops and activate ATR-Chk1 pathway where as co-directional TRCs reduce R-loops and activate ATM-Chk2 pathway (31). Head-on collisions are observed to be more deleterious and are promoted by replication stress. We reasoned that compared to normal cells, head-on collisions would be more prevalent in PDAC due to KRAS-induced replication stress. This is supported by the observation that prolonged oncogenic KRAS induction enhances R-loops (Figure 2E). We therefore hypothesized that BER inhibition by MX would enhance TRC-dependent DNA damage through ATR-Chk1 pathway as opposed to ATM-Chk2 pathway. Indeed, MX caused phosphorylation of ATR-thr1989 and Chk1-ser345 over time (Figure 5B). However, there was no change in the phosphorylation levels of ATR-ser428. Liu et. al. have previously demonstrated that ATR phosphorylation at Thr1989 (but not Ser428) is critical for Chk1 phosphorylation (38). Moreover, ATR and Chk1 phosphorylation was dependent on transcription, as co-treatment with DRB abrogated ATR-Chk1 pathway activation, both in MX treated and untreated conditions. In contrast, MX treatment did not result in ATM pathway activation and DRB did not abrogate basal levels of Chk2 phosphorylation (Figure 5C).

Collectively, these experiments demonstrate that BER pathway inhibition causes transcription-dependent DNA damage and activation of ATR-Chk1 pathway.

### BER and ATR inhibition cause synergistic toxicity in PDAC cells

Next, we hypothesized that co-inhibition of BER and ATR-Chk1 pathway would synergistically enhance DNA damage and reduce clonogenic cell survival. We analyzed DNA damage by western blot analysis of γH2AX in Panc1 and MIA Paca2 cells treated with or without MX (6-12mM) in combination with or without ATR inhibitor - VX970 (80-5000nM) for 48 hours. The results shown in Figure 5D demonstrate a slight increase in DNA damage with MX and VX970 alone compared to DMSO control in Panc 1 cells. However, there was a robust increase in DNA damage when cells were co-treated with MX (6mM) and VX970 (80nM).

To determine the consequences of DNA damage on cell proliferation, we performed a reat-time cell proliferation assay where Panc1 and MIA Paca2 cells were exposed to control conditions, single agent treatment, or a combination treatment with MX and VX970. As shown in Figure 5E and F, there was slight to no inhibition of cell proliferation with monotherapy but this was synergistically enhanced with combination treatment.

## DISCUSSION

Transcription is now recognized to be a major hinderance to replication (12). Stalled replication machinery due to TRCs generates replication stress which is exaggerated in oncogene-driven tumor cells. Here, we have a performed detailed characterization of TRCs in PDAC. Our results for the first time demonstrate that TRCs are a major feature of human PDAC.

Oncogene activation can significantly enhance protein synthesis to support the malignant phenotype (39). For instance, KRAS transformed PDAC cells consume large quantities of extracellular albumin to provide building blocks for protein synthesis (40). Oncogene-induced protein synthesis is driven by a global increase in transcription (12). Mounting evidence now supports the notion that endogenous replication stress from transcription is a major source of DNA damage in cancer cells (39). Transcription-induced DNA damage can result from prolonged stalling and collapse of replication machinery when it encounters transcription-related obstacles. These obstacles include not only the proteins involved in transcription e.g. RNAPII; but also, the transcription-induced structural changes in the DNA template e.g. topological stress or non-B DNA structures. For instance, estrogen-mediated replication stress and DNA damage in breast cancer is caused by co-transcriptional DNA:RNA hybrids (R-loops) (41). Similarly, a recent report demonstrated that oncogenic HRAS elevated the expression of general transcription factor TATA-box binding protein (12). This resulted in global increase in transcription and co-transcriptional R-loops; both of which culminated in oncogene-induced DNA damage. Consistent with published observations, we establish the causal role of oncogenic KRAS in driving TRCs, R-loops and TRC-mediated genomic instability and extend these observations to PDAC.

Emerging data demonstrate that factors in the TRC resolution pathways are critical to prevent DNA damage and loss of these factors is detrimental to cell survival. For instance, in one TRC resolution pathway, a DNA helicase (RECQL5) mediates systematic destabilization of RNAPII as the replication fork approaches transcribed chromatin (16). Loss of RECQL5 or it’s RNAPII interacting domain causes profound DNA damage through increase in TRCs. As another example, a key class of enzymes, topoisomerases, are crucial to relieve topological stress from supercoiling of the DNA. One such enzyme, topoisomerase 1 (TOP1), is critical to relax negative super-coiling at transcribed genes. Knock-down of TOP1 facilitates R-loop formation at highly transcribed genes which can stall replication forks resulting in lethal DNA damage (42). While the mechanistic insight on TOP1 function is recent, TOP1 inhibitor, irinotecan, is part of first line chemotherapy for PDAC (43). Because TOP1 function is not exclusive to TRC resolution, more specific targets in the TRC resolution pathway need to be characterized to minimize toxicity to normal cells. The current investigation, for the first time, identified a major role of BER pathway in mitigating TRCs and TRC-dependent DNA damage in PDAC.

In PDAC, BER pathway could serve as a mechanism to curtail DNA damage from oxidative stress and promote cancer cell survival. Therefore, inhibition of BER pathway could be therapeutically advantageous (44,45). Indeed, genetic or pharmacologic targeting of BER pathway proteins such as APE1 or XRCC1 has been shown to be effective in murine models of PDAC (44,46,47). The current investigation has identified a major non-canonical role of BER factors in TRC mitigation. Pharmacologic inhibition of BER pathway using MX or CRT0044876 enhanced TRCs subsequently resulting in DNA damage. Mechanistically, we demonstrate that the major impact of BER pathway inhibition is at the transcribed chromatin where TRCs are promoted by RNAPII stalling. Previous studies have demonstrated that guanine is the most vulnerable of the nucleobases to oxidation and leads to transcription-blocking lesions: 5-guanidinohydantoin (Gh) and spiroiminodihydantoin (Sp) (48). Recent structural studies have delineated a helix distorting mechanism of RNAPII stalling at these DNA lesions (49). Subsequently, we demonstrate that both replication and transcription machinery are dislodged from transcriptionally active chromatin resulting in DNA damage.

We demonstrate that BER inhibition-related DNA damage triggers predominantly ATR-Chk1 pathway activation. Experiments performed by Hamperl et. al. have previously demonstrated that conflict orientation is a major determinant of checkpoint pathway activation (31). Head-on collisions predominantly activate the ATR-Chk1 pathway. Whereas co-directional collisions trigger ATM-Chk2 pathway. They further demonstrate that replication stress can increase the frequency of head-on collisions. While we did not examine conflict orientation in the present study, the robust transcription-dependent activation ATR-Chk1 pathway suggests that BER pathway may potentiate head-on TRCs. Interestingly, we find that both DNA damage and ATR-Chk1 activation can be robustly inhibited by transcription inhibitor – DRB, suggesting that BER targeting induced genotoxicity is in part transcription-dependent. These mechanistic observations uncover a potential therapeutic vulnerability in human PDAC.

While BER pathway inhibition with MX at clinically relevant dosages causes DNA damage, this led to a modest decrease in cell proliferation for both Panc1 and MIA Paca2 cells. This is likely explained by the activation of ATR-Chk1 pathway that may facilitate repair of MX-induced DNA damage. Expectedly, inhibition of ATR using VX970 and BER pathway using MX was synergistic in producing DNA damage. Likewise, the combination was synergistic in inhibiting cell proliferation in both Panc1 and MIA Paca2 cells. Both MX and VX970 have been used in early phase clinical trials but not in combination (50,51). These findings provide a mechanistic rationale for further pre-clinical development of BER pathway inhibitors in combination with ATR pathway inhibitors.

## Supporting information

Supplemental Results

## ACKNOWLEDGEMENTS

Research reported in this publication included work performed in the Analytical Cytometry and Light Microscopy & Digital Imaging Shared Resource supported by the National Cancer Institute of the National Institutes of Health under grant number P30CA033572. The content is solely the responsibility of the authors and does not necessarily represent the official views of the National Institutes of Health. This work was funded by the NCCN Foundation®. Any opinions, findings, and conclusion expressed in this material are those of the author(s) and do not necessarily reflect those of National Comprehensive Cancer Network® (NCCN®) or the NCCN Foundation.

