## Supplemental Results for "Base Excision Repair Pathway Regulates Transcription-Replication Conflicts in Pancreatic Ductal Adenocarcinoma"

Fan Meng, Anup K. Singh, Tiane Li, Marc Attiyeh, Fatemeh Kohram, Terence Williams, Yilun Liu, Mustafa Raoof
City of Hope Cancer Center, Duarte, CA 91010

**Figure S1**

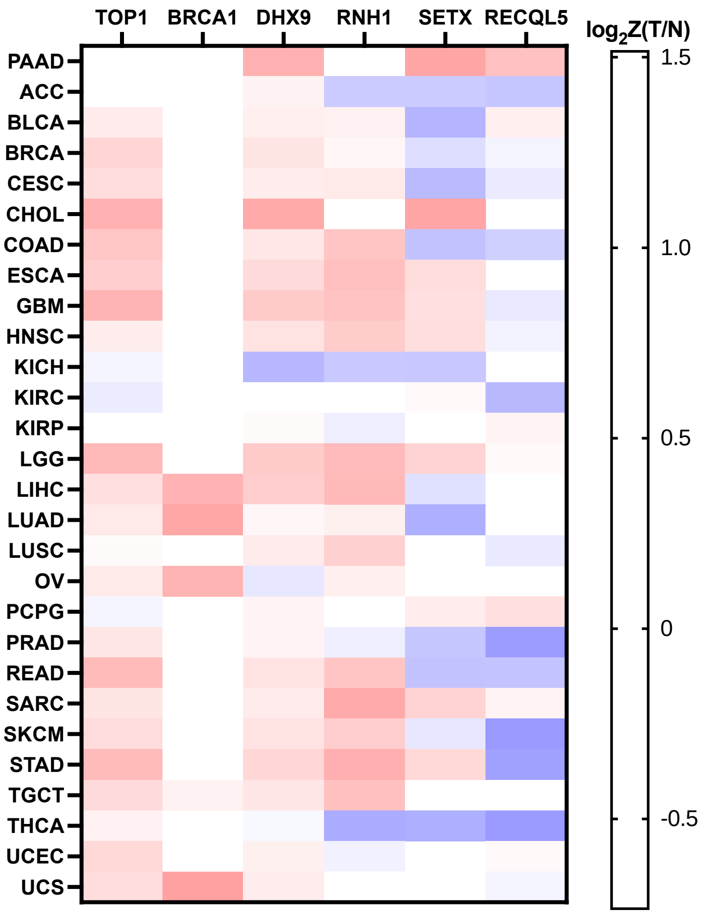


**Figure S1. Human PDAC demonstrates high levels of TRCs**

**Gene-expression analysis** of known proteins involved in TRC resolution pathways are uniquely upregulated in PDAC tumors (**T**) vs normal tissue (**N**) in comparison to other tumor types. (Tumor type abbreviations: **PAAD**-Pancreatic adenocarcinoma, **ACC**-Adrenocortical carcinoma, **BLCA**-Bladder Urothelial Carcinoma, **BRCA**-Breast invasive carcinoma, **CESC**-Cervical squamous cell carcinoma and endocervical adenocarcinoma, **CHOL**-Cholangiocarcinoma, **COAD**-Colon adenocarcinoma, **ESCA**-Esophageal carcinoma, **GBM**-Glioblastoma multiforme, **HNSC**-Head and Neck squamous cell carcinoma, **KICH**-Kidney Chromophobe, **KIRC**-Kidney renal clear cell carcinoma, **KIRP**-Kidney renal papillary cell carcinoma, **LGG-**Brain Lower Grade Glioma, **LIHC**-Liver hepatocellular carcinoma, **LUAD**-Lung adenocarcinoma, **LUSC**-Lung squamous cell carcinoma, **OV**-Ovarian serous cystadenocarcinoma, **PCPG**-Pheochromocytoma and Paraganglioma, **PRAD**-Prostate adenocarcinoma, **READ**-Rectum adenocarcinoma, **SARC**-Sarcoma, **SKCM**-Skin Cutaneous Melanoma, **STAD**-Stomach adenocarcinoma, **TGCT**-Testicular Germ Cell Tumors, **THYM**-Thymoma, **THCA**-Thyroid carcinoma, **UCEC**-Uterine Corpus Endometrial Carcinoma, **UCS**-Uterine Carcinosarcoma)

Figure S2

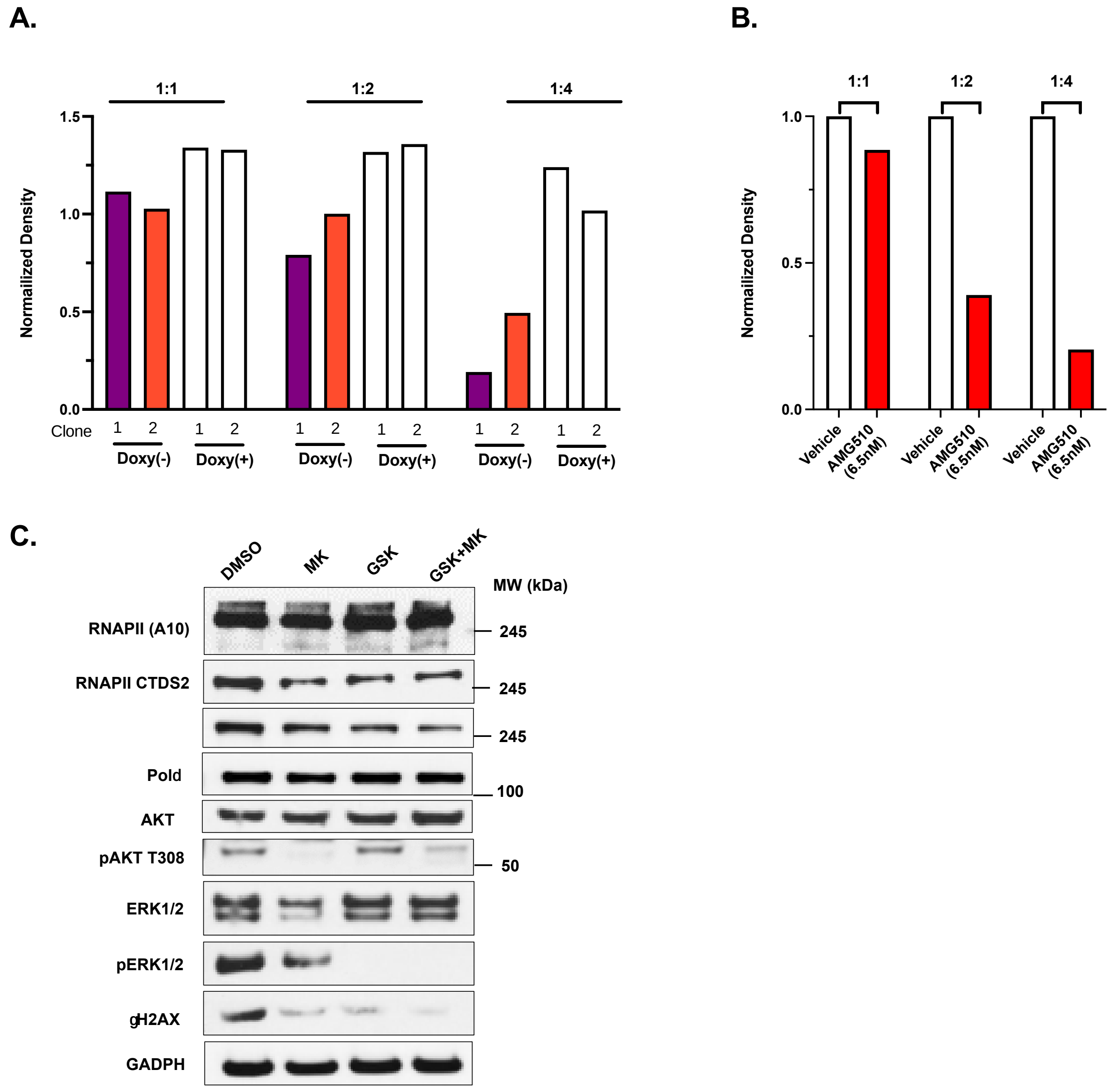


**Figure S2. Oncogenic KRAS causes TRCs and DNA damage**

A. **Dot blot assay.** Densitometry quantification of blot shown in Figure **2E**. Values are normalized to parental control.

**B. Dot blot assay.** Densitometry quantification of blot shown in Figure **2F**. Values are normalized to Vehicle control for each dilution.

**C. Western blot.** Representative Western Blot analysis of MEK and AKT pathway signaling in Panc1 whole-cell extracts prepared from control (DMSO), MEK inhibitor (GSK1120212, GSK) or AKT inhibitor (MK2206, MK)-treated cells as indicated. GADPH was used as a loading control.

**Figure S3**

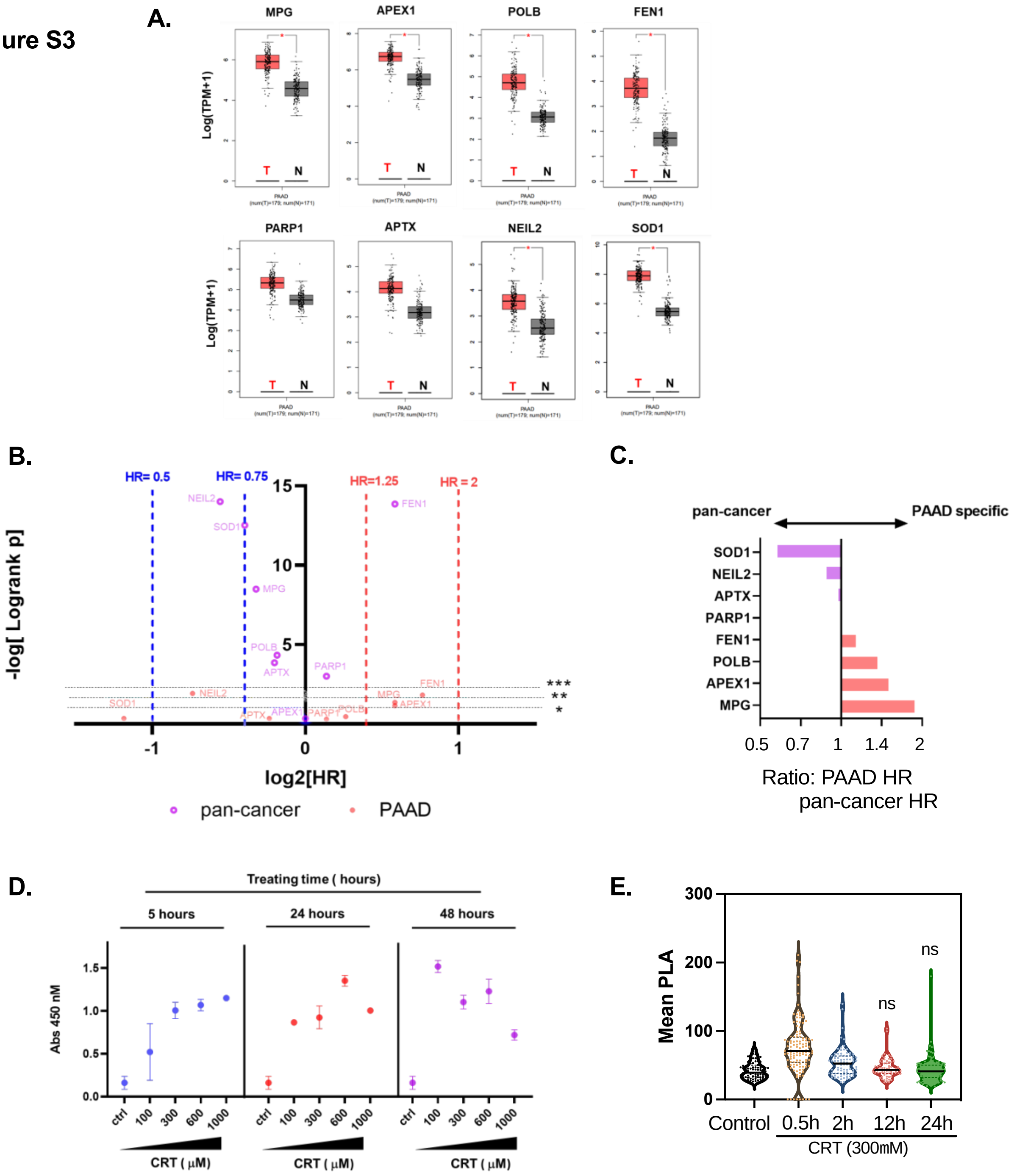


| 5 hours CRT treating | | 24 hours CRT treating | | 48 hours CRT treating | |
|---|---|---|---|---|---|
| CRT ( μM) | AP sites/100 kbp | CRT ( μM) | AP sites/100 kbp | CRT ( μM) | AP sites/100 kbp |
| Ctrl | 3 | | | | |
| 100 | 12 | 100 | 33 | 100 | 36 |
| 300 | 23 | 300 | 22 | 300 | 26 |
| 600 | 25 | 600 | 32 | 600 | 29 |
| 1000 | 31 | 1000 | 23 | 1000 | 16 |

**Figure S3. BER pathway regulates TRCs in PDAC cells**

**A. Gene-expression analysis** of TRC-associated BER proteins identified in Figure 3B are upregulated in PDAC tumors (**T**) vs normal tissue (**N**) (All T vs. N comparisons – Student t-test, p-value <0.001). Tumor data source TCGA project, Normal data source is GETx project.

**B.** Association of gene expression of TRC-associated BER proteins and overall survival in Pancreatic Cancer (PAAD) and pan – cancer. For each gene, gene expression was categorized as high or low at a median cut-off level. (P-value was calculated using unadjusted Logrank test).

**C.** Ratio of PAAD hazard ratio vs. pan cancer hazard ration of death. Most TRC-associated BER proteins demonstrated worse survival in PAAD vs. pan-cancer (high ratio).

**D. Detection of apurinic/apyrimidinic (AP) sites upong exposure to APE1 inhibitor (CRT0044876).** Panc1 cells were treated with increasing concentrations of CRT or control; and AP sites were quantified using a commercial colorimetric assay kit (Cell BioLabs, STA-324).

**E. Immunocytochemistry.** RNAPII-PCNA PLA was performed to quantify TRCs after indicated treatment conditions with CRT. Representative experiment is shown. (****p<0.0001, Welch's unpaired t-test vs. control). CRT=CRT0044876.

**Figure S4**

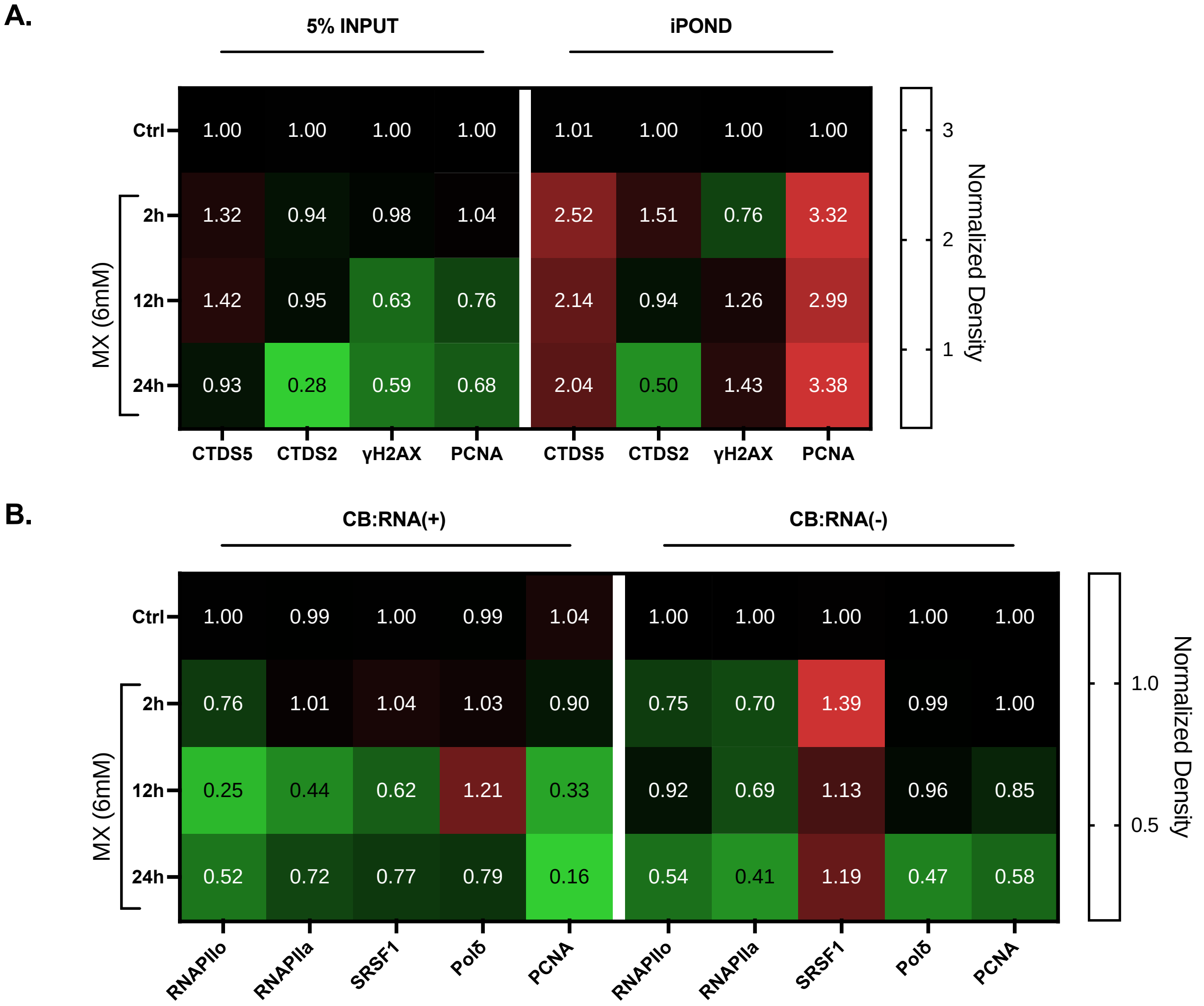


**Figure S4. BER inhibition impacts RNAPII dynamics at TRCs.**
**A.** Densitometry Quantification of Blots shown in Figure 4A. The densitometry values are first normalized to the loading control (Histone H3) and then normalized to control (no treatment).
**B.** Densitometry Quantification of Blots shown in Figure 4Bii. The densitometry values are first normalized to the loading control (Lamin A/C) and then normalized to control (no treatment).
